# Leveraging Persistent Homology of Eye Movements for Neural Disorder Screening

**DOI:** 10.1101/2024.11.23.624966

**Authors:** Dongcheng He, Shaoying Wang, Haluk Ogmen

**Author notes:** Corresponding Author: Dongcheng He, 288 Chuangxin Blvd, High-Tech District, Hefei, Anhui, China, 230071.

## Abstract

Abnormal ocular behaviors are associated with numerous neural disorders, detectable through specific eye movement patterns. Eye tracking—a non-invasive, accessible method—has thus been investigated as a tool for the automated diagnosis of these disorders. However, traditional eye movement feature extraction methods have limitations and are not universally applicable across diverse tasks. In this study, we present a novel feature extraction approach using Persistent Homology, a topological data analysis technique, to capture spatial and temporal topological features from eye movement data. Topological features retain multiple theoretical properties, making them robust for identifying abnormal eye movements and broadly applicable across tasks. By using only spatio-temporal topological features, our approach demonstrated promising performance in screening dyslexia and ADHD with a simple shallow perceptron classifier containing one or two hidden layers. These results support the informative validity of topological features as well as the efficacy of our proposed methods for extracting them.

## 1. Introduction

Eye movements exhibit complex behaviors driven by various cortical systems, such as oculomotor control and spatial attention, distributed across various brain regions (Leigh & Lee, 2015). Consequently, abnormal eye movements may reflect impairments that range from mechanical to more complex neurological and psychological causes with patterns varying based on the disorder type and specific task requirements (Anderson & MacAskill, 2013; Kuskowski, 2013; Rommelse et al., 2008; Armstrong & Olatunji, 2012). These abnormalities provide critical insights into disrupted neural processes and pathways underlying each condition. For instance, studies have shown that individuals with ADHD often exhibit unstable fixational and saccadic eye movements, largely due to reduced inhibitory control associated with frontal lobes (Munoz et al., 1999; Munoz et al., 2013; Goto et al., 2010). Disorganized visual scanning is frequently observed in patients with Alzheimer’s disease, especially during reading tasks, associated with degeneration in the frontal cortex (Lueck et al., 2000; Fernández et al., 2013; Parkinson & Maxner, 2005; Fletcher & Sharpe, 1986). Reduced fixation stability in amblyopic eyes has been found to be primarily limited by microsaccade errors (Chung et al., 2015). These characteristics underscore the potential of eye-tracking as a valuable tool for neural disorder screening.

Training machine learning models on eye-tracking data to detect neurological and cognitive impairments has been actively explored. Studies have demonstrated that eye-movement data, either independently or combined with multimodal data, can effectively predict disorders such as Parkinson’s disease (Przybyszewski et al., 2014; Tseng et al., 2013), Alzheimer’s disease (Jang et al., 2021), autism (Vabalas et al., 2020; Meng et al., 2023), ADHD (Deng et al., 2022; Yoo et al., 2024; Tseng et al., 2013; Lee et al., 2023), and dyslexia (Raatikainen et al., 2021; Nilsson Benfatto et al., 2016), achieving promising accuracy against healthy controls. Although these conditions induce distinct ocular behaviors across various tasks, feature extraction from eye movements typically follows a similar protocol. Due to the high-dimensional, event-related, time-series nature of lab-collected eye-movement data, the analysis of these data often involves segmenting critical event-windows within each sample and quantifying them using metrics such as fixation duration and saccade orientation (Nilsson Benfatto et al., 2016; Zemblys et al., 2018). This approach presents two major limitations for the use of abnormality screening: First, reducing noise in these features requires simplified experimental conditions; second, it overlooks broader patterns within the eye-movement data. Previous studies have used multimodal data to overcome these limitations. For instance, Tseng et al. (2013) used both eye-movement data and corresponding stimulus images as inputs into deep learning models to enable the extraction of higher-level features (Tseng et al., 2013). However, the gap in developing an effective feature-extraction method generally applicable to a broad range of tasks has not been filled.

In this study, we propose a novel feature-extraction method using Topological Data Analysis (TDA) applied to eye-movement data. TDA is an emerging framework for analyzing large-scale data through geometric and algebraic topology methods (Carlsson, 2014). It operates on a simplicial complex, a mathematical construct that describes a set of simple shapes (e.g., points, lines, triangles) connected in a specific way to form more complex structures or spaces. By applying TDA, we can capture topological features of various dimensions, known as homology, in the simplicial complex. For example, 0-order homology represents connected components, while 1-order homology represents loop patterns, and so on. When eye-movement data, such as gaze trajectories sampled at specific intervals, are represented as a point cloud in Euclidean space, a simplicial complex can be defined using the Vietoris–Rips approach (Vietoris, 1927; Chambers et al., 2010; Carlsson, 2014). In this approach, points serve as vertices, and an edge is established between two vertices if the distance between them falls below a given threshold. By adjusting this threshold, the point cloud data reveals different patterns of simplicial complex structures.

A central technique in TDA is persistent homology (PH), which captures the topological structure of a finite point set by observing the appearance and disappearance of homological features across varying threshold values (Robins, 1999). The threshold values at which a topological feature appears and then disappears are known as its “birth” and “death”, respectively, with the difference referred to as its “lifetime”. Mapping birth values on the x-axis and death values on the y-axis, a persistence diagram can be constructed to visualize all topological features in a single chart, which provides a straightforward summary of the topological properties stored in the data.

This approach offers several advantages that address the limitations of traditional feature extraction methods from eye-movement data. First, PH distinguishes short-lived features, likely attributable to noise, from long-lived features that remain stable across multiple scales, demonstrating high noise-tolerance. Second, homological features are preserved under transformations such as scaling and rotation, allowing fundamental characteristics to be identified regardless of variations in orientation or shape. Third, PH enables the extraction of shared features across individuals, facilitating the identification of common behaviors within specific populations while minimizing information loss, as it does not rely on assumptions like Gaussian distributions. Finally, this method effectively reduces high-dimensional data to simpler topological representations, providing a structured summary of the data’s global shape. These properties make PH an effective tool for analyzing eye movement data that may contain unstable ocular motor behaviors, especially in complex scenarios like viewing natural scenes. Additionally, by reducing rather than increasing data dimensionality, this approach is well-suited for machine learning applications.

In our method, we projected spatial and temporal patterns of eye movements into the Euclidean space and performed persistent homology (PH) on each dimension independently. Using the resulting topological features, we conducted two neural-disorder screening tasks: dyslexia and ADHD. Our results suggest that this approach can effectively capture topological features for abnormality detection and might be broadly applicable to general eye-movement analysis, as demonstrated by promising performance in our experiments.

## 2. Experiment 1

### 2.1. Methods

The data for this experiment was curated from an open-access dataset comprising eye movement recordings from 185 subjects (88 with dyslexia, 97 healthy controls) aged 8-9 years, with IRB approvals in place (Benfatto et al., 2016 a, b.). Eye movements were recorded using a goggle-based infrared corneal reflection system, the Ober-2™ (formerly Permobil Meditech, Inc., Woburn, MA), at a sampling rate of 100 Hz. During data collection, each subject read an age-appropriate passage consisting of 10 sentences across 8 lines. Subjects’ heads were stabilized using a chin rest positioned 45 cm from the display, and calibration was performed before recording. The average time for the task was 17.47±3.19 seconds, resulting in a data size of 1747±319 samples. For the analysis, only left-eye movement data was used.

Our data processing was performed using a Python (Version: 3.11.7) environment in Jupyter Notebook (Version: 7.0.8). We began by mapping the gaze trajectory onto a two-dimensional space, disregarding time. Using Giotto-tda (Version: 0.3.1, Tauzin et al., 2021), a Python based package, we performed PH on each sample and extracted the lifetimes of all 0- and 1-order homology within the spatial domain. Next, we used time-delay embedding to project the temporal dynamics of eye movements of each subject along the horizontal and vertical axes into Euclidean space using the following equation:

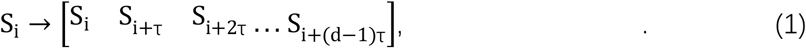

where S_i_ is a time series, and it is projected to a vector of d-dimensions with a window size τ.

In this experiment, τ was set to 1 and d was 2.

To illustrate how this procedure benefits feature extraction of temporal dynamics using TDA, we provide a simple example comparing a periodic time series with a random time series, displaying their respective projections. Figure 1 shows the projections of three time series, each of length 100. Two series are defined by periodic functions: Equation 2, a sine function with k equal to 10 and 30 respectively, while the third is a random sequence with values ranging between -1 and 1. As shown in Figure 1, regular loops can be observed for the periodic functions with diverse shapes due to different temporal frequencies, whereas random sequence is transformed to distributed scatters with irregular consisting of more complex topological features. The results demonstrate distinct topological structures highlighting the approach’s sensitivity to underlying temporal patterns.

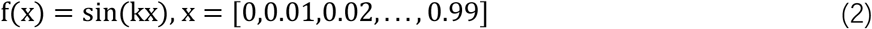

**Figure 1.**
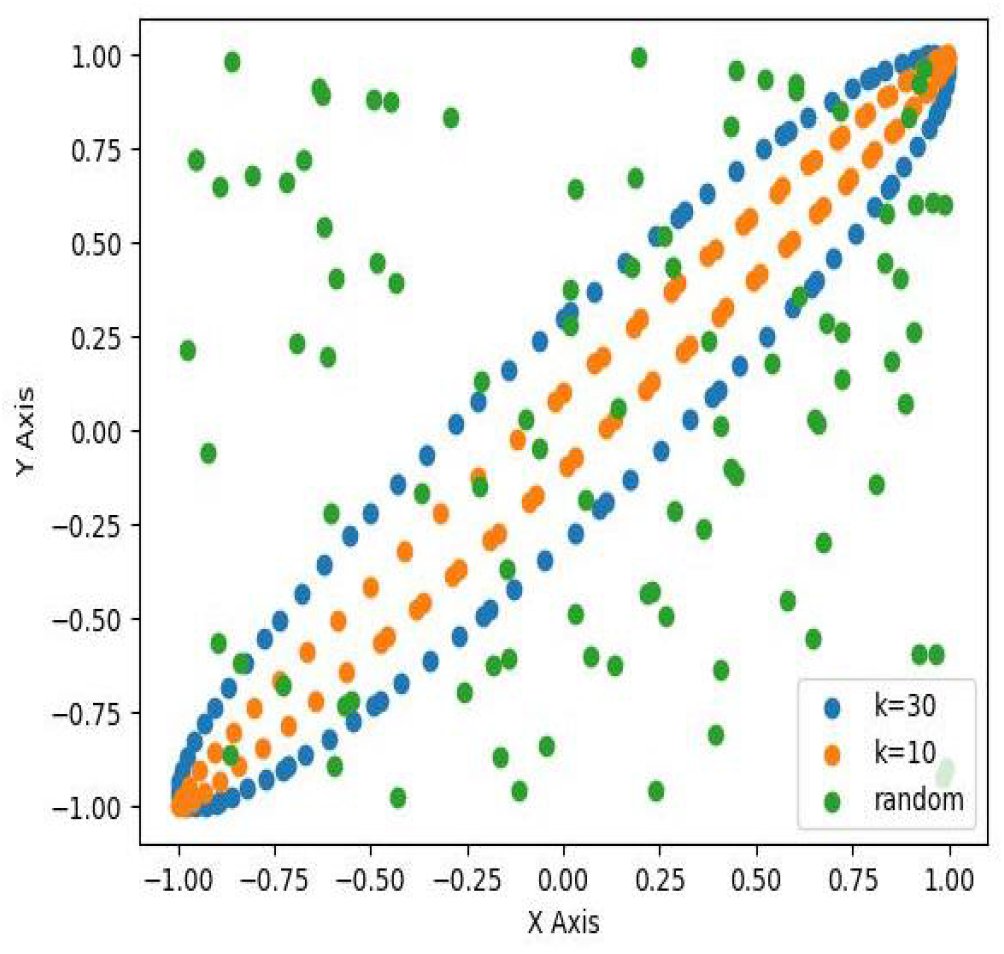
The projections of a random and two periodic time-series via time-delay embedding. Periodic time-series are sinusoidal functions defined in Eqn. (2).

After projecting the horizontal and vertical components of eye movements into the Euclidean space, through a similar PH approach, we extracted the lifetimes of all 0- and 1-order homology of eye movements from the temporal domain. Subsequently, we calculated the mean and standard deviation (two measurements) of the birth time and lifetime (two properties) of the topological features across each order and domain to represent the eye movement data. These procedures resulted in a total of 24 features, including 8 features in the spatial domain (2 orders × 2 properties × 2 measurements) and 16 features in the temporal domain (2 orders × 2 properties × 2 measurements × 2 axes). Following feature extraction, principal component analysis (PCA) was applied, and the components explaining 95% of the variance were fed into a three-layer perceptron classifier, with 4 neurons in the middle layer, to classify subjects as either with or without the dyslexia. The data was randomly split into five folds for training and testing (4:1 ratio), with this process repeated 20 times using different random sequences to ensure robustness in the results. Performance is evaluated using the metrics of Receiver Operating Characteristic (ROC) and the Area Under Curve (AUC).

### 2.2. Results

To illustrate the outcomes of each processing step, we used data from one subject as an example. Figure 2 shows the spatial trajectory of the subject’s eye movements during the reading task, sampled at 100 Hz. The resulting persistence diagram from PH analysis of the eye movement trajectory is presented in Figure 3, capturing both 0- and 1-order features through their “birth” and “death” threshold values.

**Figure 2.**
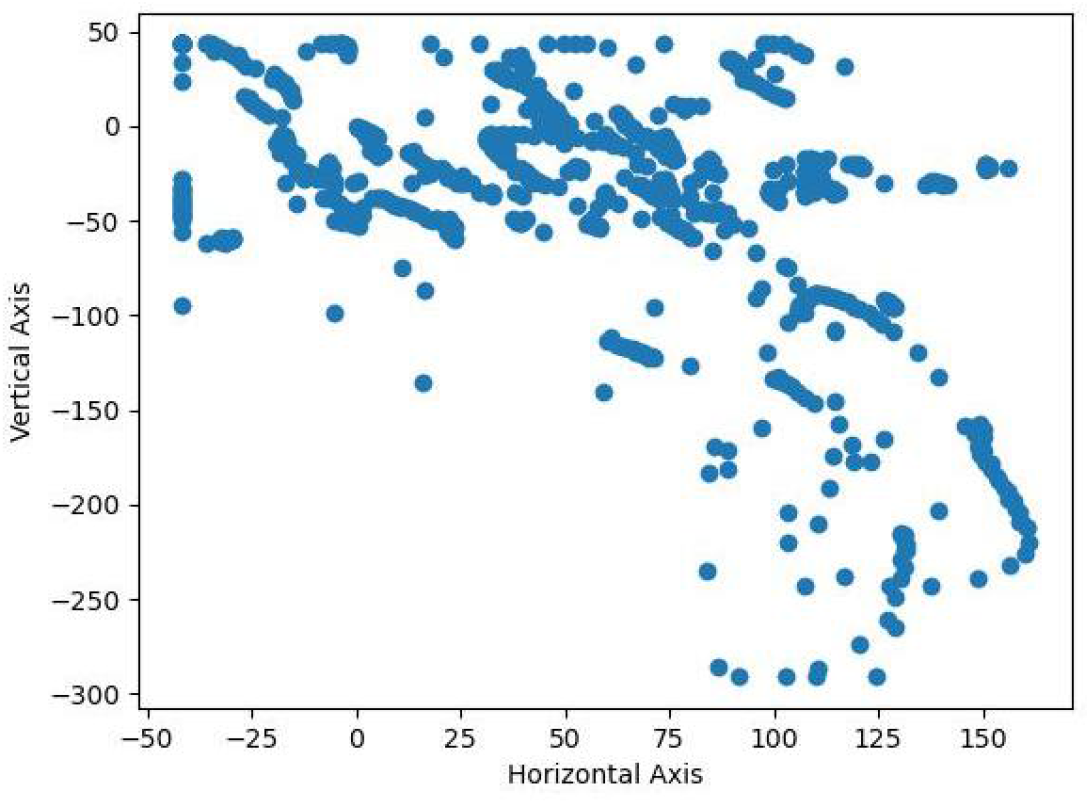
An example of subject’s eye movement trajectory during the reading task.

**Figure 3.**
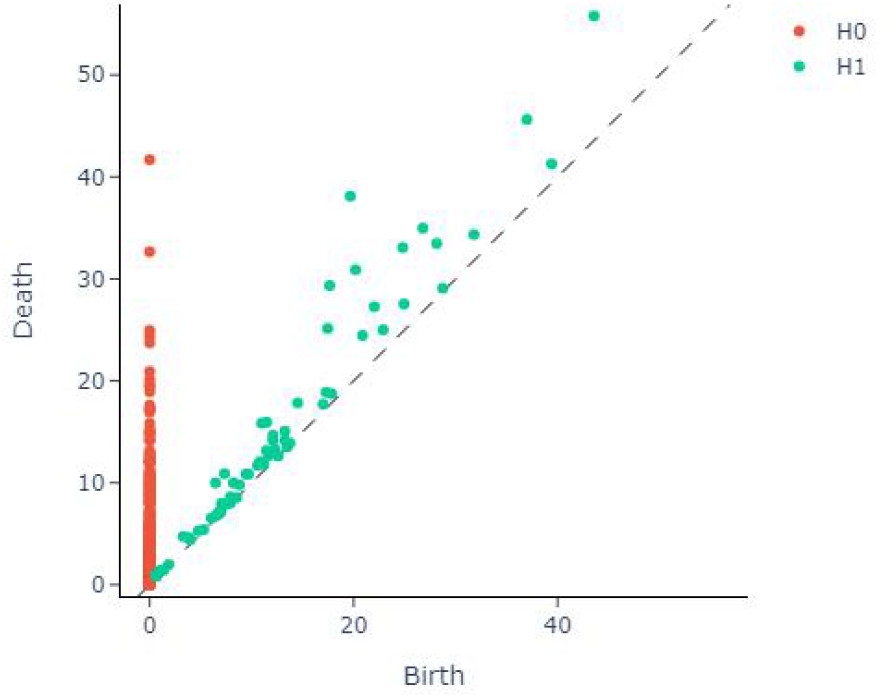
The persistence diagram given the topological structures shown in Figure 2. Each dot represents a unique topological feature. H0 and H1 indicate 0-order and 1-order respectively. The X-axis indicates the “birth” thresholds, while the Y-axis indicates the “death” thresholds. The dashed line marks the boundary where death and birth thresholds are equal, leaving any dot’s distance to which equates its “lifetime”. All persistence diagrams plotted in this paper are presented in the same manner.

After applying time-delay embedding, the projected data for the horizontal and vertical components of the trajectory are shown in Figure 4. The corresponding persistence diagrams for each component, produced by PH analysis, are displayed in Figure 5.

**Figure 4.**
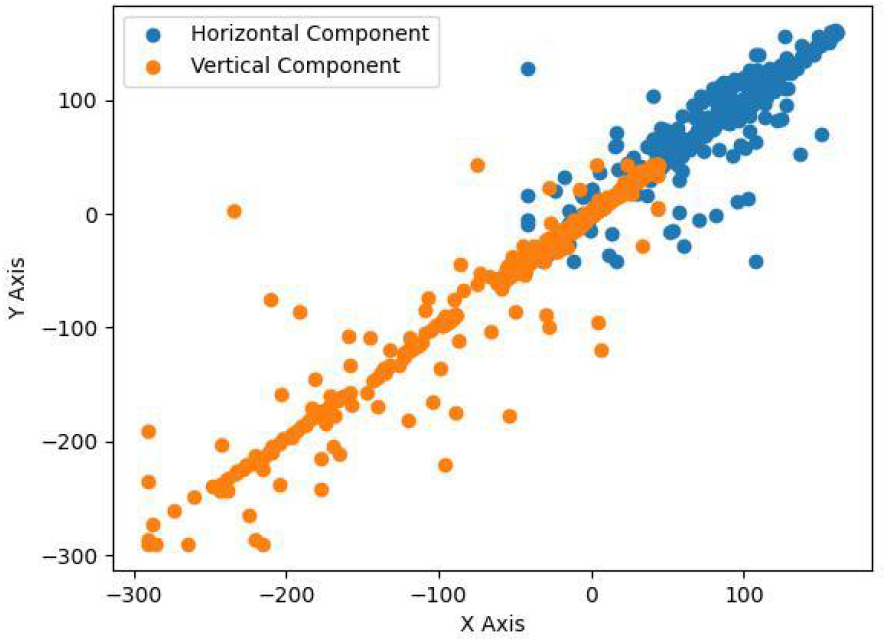
The projected horizontal and vertical components of a subject’s eye movements via time-delay embedding.

**Figure 5.**
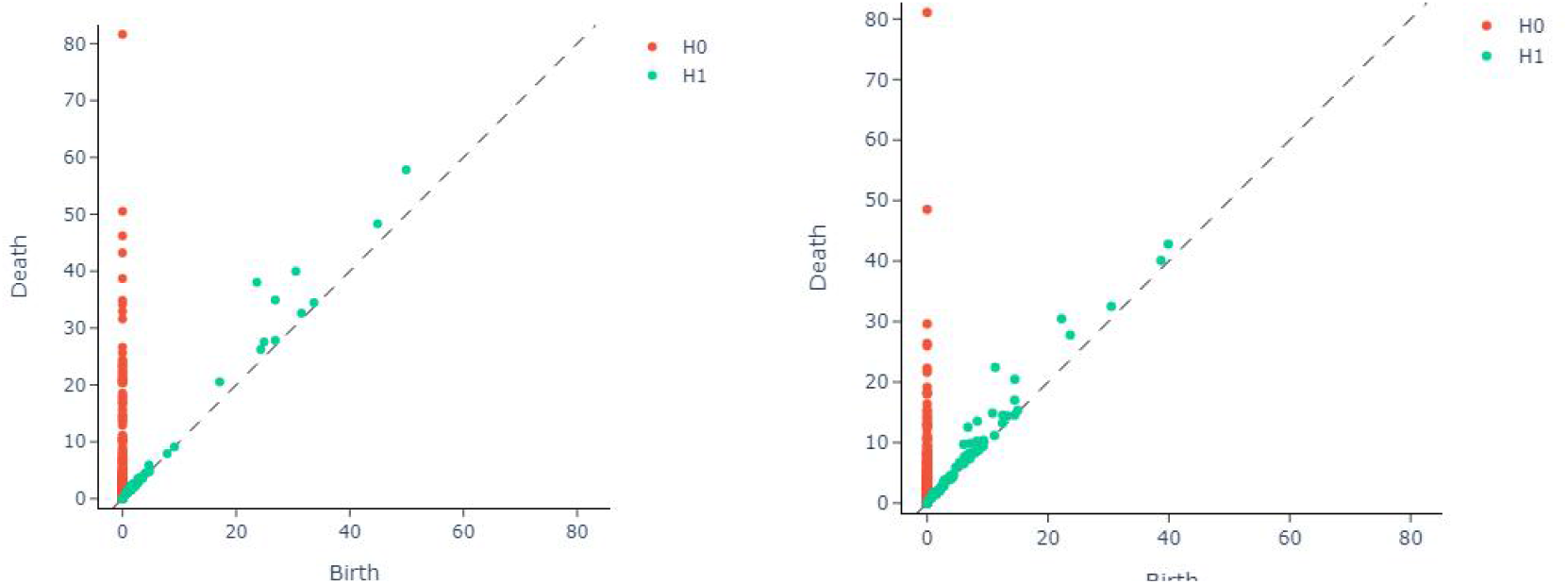
The persistence diagram given the topological structures shown in Figure 4. Left: horizontal component. Right: vertical component.

For the machine learning task, we next illustrate the feature extraction and classification results. The PCA results, shown in Figure 6, indicate that the top five principal components together explain 95.5% of the variance. Using these features, our method achieved a mean AUC of 0.91± 0.04. The ROC curves for all tests are plotted in Figure 7. Our approach demonstrated promising performance in dyslexia screening, with a simple three-layer perceptron model, using only four neurons in the hidden layer.

**Figure 6.**
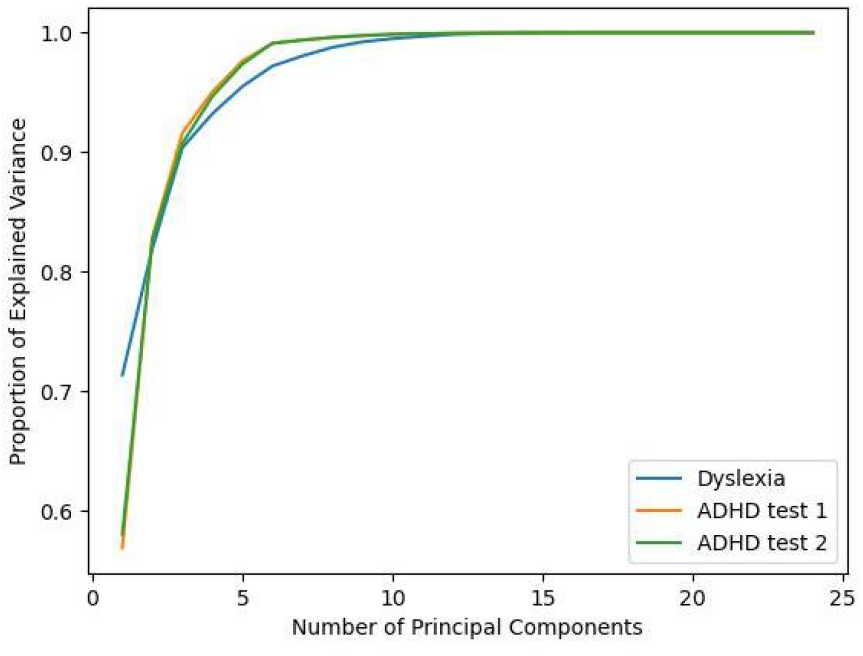
Results from principal component analysis (PCA).

**Figure 7.**
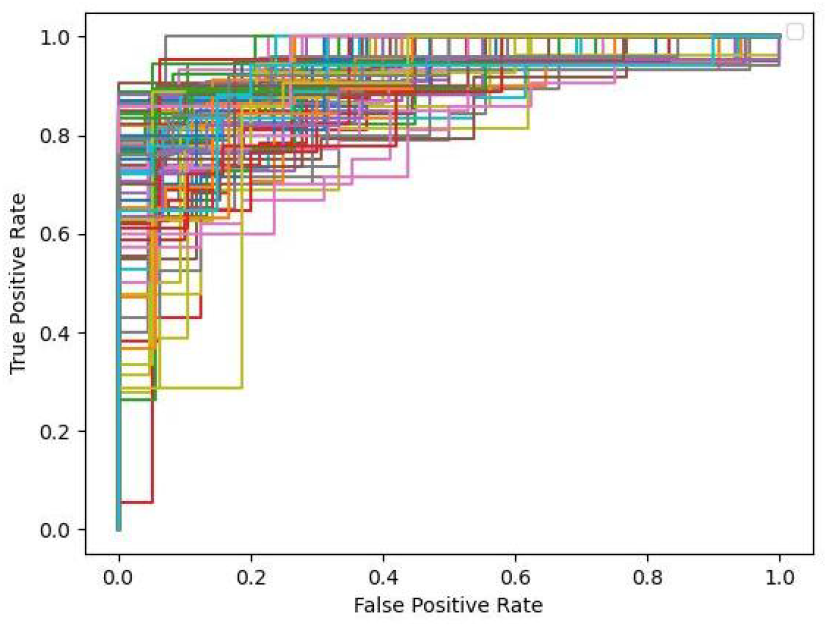
ROC curves for all the testing sessions in the Experiment 1. Each color represents a unique session of test.

## 3. Experiment 2

### 3.1. Methods

The data for this experiment was curated from an open-access dataset containing eye movement records of 50 subjects (28 with ADHD, age 10.71±0.54 years; 22 healthy controls, age 11.58±0.5 years) collected during a sequential cognitive task, with IRB approvals (Rojas-Líbano et al., 2019 a, b.). Eye movements were recorded using the Eyelink 1000 system (SR Research Ltd., Mississauga, Ontario, Canada) at a sampling rate of 1000 Hz. Subjects were seated at a table equipped with a computer screen for task presentation and the eye-tracking device, with their heads stabilized using a forehead/chin rest (SR Research Ltd.) positioned 60 cm from the screen. For ADHD patients, both on- and off-medication conditions were tested, though only the off-medication data were included in this experiment.

Each task session in this experiment comprised eight consecutive blocks, each containing a sequence of 20 trials (schematic shown in Figure 8). During each trial, subjects observed a sequence of three dot arrays presented on a 4×4 grid, separated by a 500ms fixation period. After the dot sequences, a distractor image was shown; this distractor could be a dot array, an emotional image, a neutral image, or a flat-color screen. Following the distractor, a probe dot array was presented, and the subject had to indicate whether the probe dot appeared in any of the previous three arrays. After 1.5 seconds, subjects received feedback on their performance for the trial.

**Figure 8.**
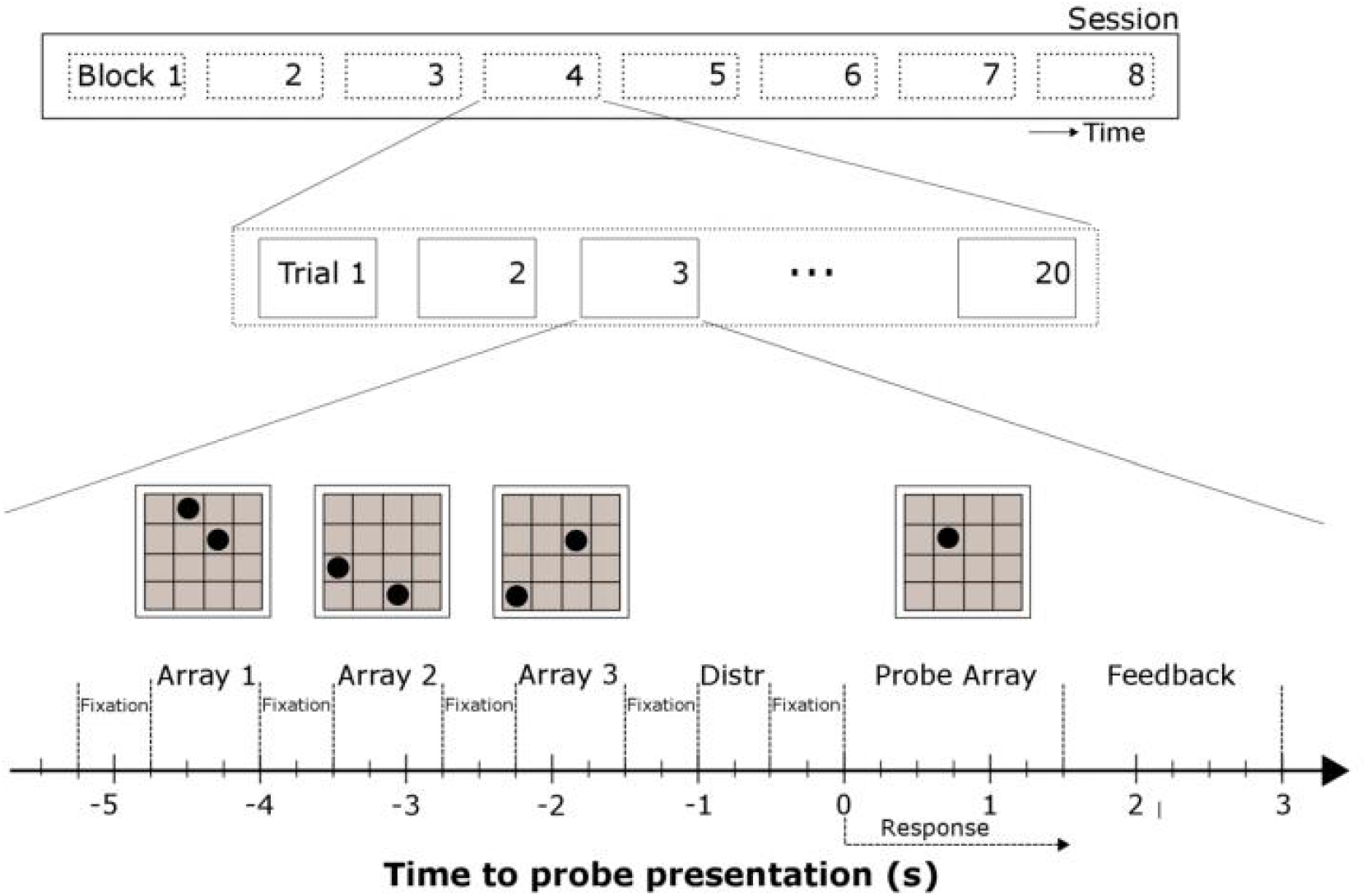
Schematics of the experimental procedures for the dataset used in Experiment 2. This figure was captured from the Fig.2 of Rojas-Líbano et al., 2019 a, the dataset descriptor.

In this experiment, we conducted two tests with different data corresponding to different stimuli. In this first test, only the data related to single-dot arrays was used in our study, as our preliminary analysis indicated that this stimulus resulted in the best performance. Each dot array was displayed for 750 ms, resulting in a data length of 750 points per trial. Trials in which subjects closed their eyes for more than 100 ms were excluded based on pupil size monitoring. This led to an average of 202.22±34.72 trials for ADHD subjects and 224.48±17.82 trials for healthy controls. The total number of samples analyzed was 5,460 for ADHD subjects and 4,714 for healthy controls. In the second test, we include the data correspondent to all six types of stimuli during the array sequence and distractor presentations. This led to an average of 546.77 ±90 trials for ADHD subjects and 603.29±53.13 trials for healthy controls. The total number of samples analyzed was 14,763 for ADHD subjects and 12,669 for healthy controls.

The data processing procedures for this experiment were the same as in Experiment 1. Following feature extraction, principal component analysis (PCA) was applied, and the components explaining 99% of the variance were fed into a perceptron classifier. In the first test, with a relative small data size, we used a three-layer perceptron with 20 neurons in the hidden layer to classify a trial as either with or without ADHD. The data were randomly split into ten folds for training and testing (9:1 ratio), with this process being repeated 10 times using different random sequences to ensure robustness in the results. In the second test, a four-layer perceptron with 80 and 40 neurons in the two hidden layers was used. The data were randomly split into ten folds for training and testing (9:1 ratio) for once.

### 3.2. Results

The PCA results for both tests are plotted in Figure 6. Seventeen principal components were used in both tests. In the first test, the mean AUC of ROC was 0.66±0.01. In the second test, the mean AUC of ROC was 0.69±0.01. ROC curves for both tests are shown in Figure 9. We found that our approach achieved promising performance in screening ADHD using simple perceptron models and eye movement data with a time duration no longer than 750ms.

**Figure 9.**
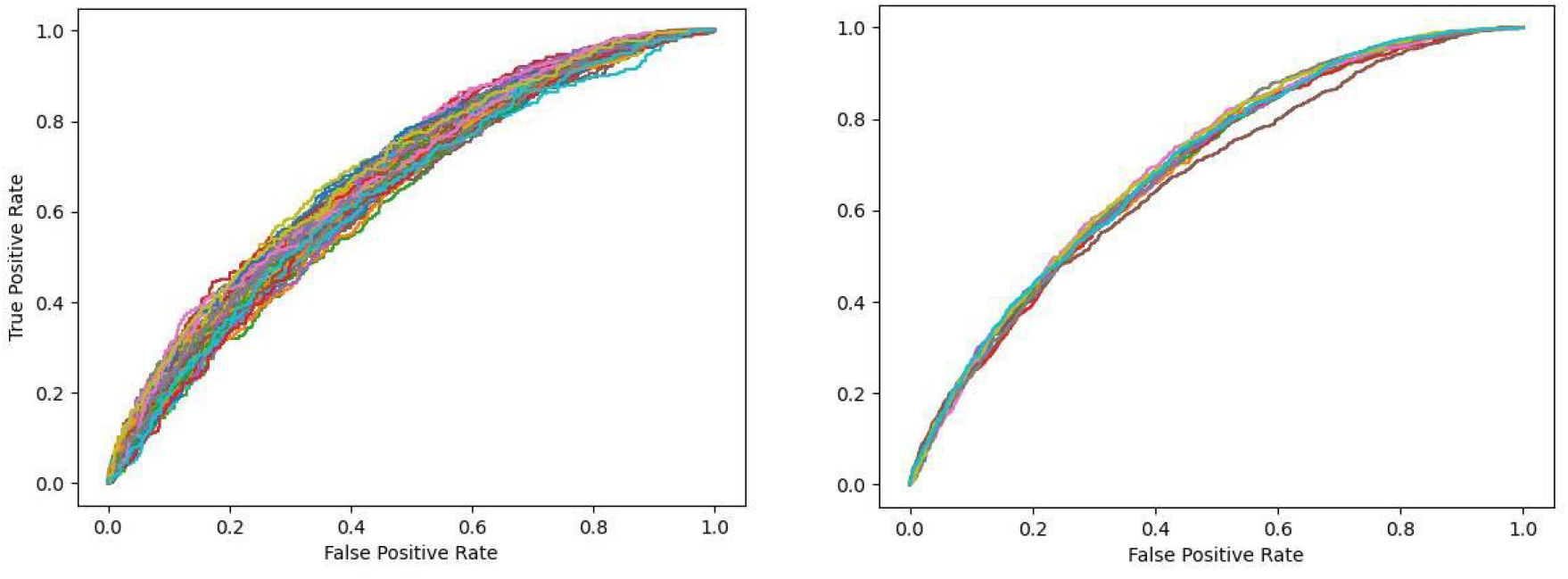
ROC curves for all the testing sessions in the Experiment 2. Left: test 1 with one-dot array related data. Right: test 2 with six types of stimuli.

## 4. Discussion

Eye tracking as a non-invasive diagnostic tool for neural disorders by identifying distinct eye movement patterns associated with specific conditions has been actively explored. Given the recent advances in developing cheaper and more flexible eye-trackers, the utility of these approaches is expected to increase. Traditional methods of feature extraction from eye-movement data have inherent limitations, which our study addresses through the introduction of a novel approach based on persistent homology, a method of topological data analysis. By capturing complex topological features across both spatial and temporal domains, our method shows substantial potential in improving classification for disorders, as proved by our pioneering tests in predicting dyslexia and ADHD.

Without a standard dataset for model validation and given the limited published results, it is challenging to perform a comprehensive comparison of our performance with that of previous studies. In this section, we discuss several recent studies that utilized eye-movement data exclusively, without integrating features from other modalities. For instance, Benfatto et al. (2016) evaluated over 150 features derived from eye-movement data on a reading task and observed a positive correlation between the number of features and classification accuracy, achieving a maximum accuracy of 95.3%. In a separate study, Raatikainen et al. (2021) employed a support vector machine (SVM) model to classify dyslexia, attaining a best accuracy of 89.7% with a recall of 84.8%. Regarding ADHD, Deng et al. (2022) analyzed eye movement data from a video-watching task, using a convolutional neural network (CNN) to differentiate ADHD patients from healthy controls, achieving a mean AUC of 0.646. Additionally, Yoo et al. (2024) collected eye movement data during behavioral tasks assessing working memory and attentional effort in ADHD and healthy subjects, reaching an AUC of 0.66 using only eye tracking data. Although these studies employed distinct experimental setups during data collection, these findings indicate that our results are comparable for dyslexia and ADHD screening tasks.

More importantly, the main aim of this study was to propose a feature-extraction method that is potentially generally applicable to a broad range of tasks using eye-tracking data, especially neural disorder screening. In our experiments, we used the same methods in both dyslexia and ADHD screening tasks, without any distinction, and achieved promising performance in both tasks. As a parameter-free method, model performance is more robust for interpretation. In addition, to explicitly examine the informative validity of the topological features, we adopted simplified models (perceptron classifiers with one or two hidden layers) and feature selection methods (statistical abstraction of topological features). Our test results provide strong evidence that topological features extracted by persistent homology are informative in differentiating abnormal eye movements.

It should be acknowledged that the topological patterns captured in our feature set likely represent only a subset of the potential information embedded in these features. Due to constraints in data size and an unalterable experimental setup, our study was limited in its ability to perform more comprehensive feature engineering on the homology patterns. For future work, we recommend a more extensive extraction of patterns from spatio-temporal topological features, such as calculating second-order homology by using three-dimensional eye-tracking data combined with three-dimensional time-delay embedding. This approach could reveal richer topological structures, potentially enhancing model performance and interpretability in neural disorder classification.

## Acknowledgments

The authors declared no funding source in support of this study and no conflict to interest.

## Author Contribution

DH: Conceptualization, Data curation, Formal analysis, Investigation, Methodology, Project administration, Resources, Software, Validation, Visualization, Writing-original draft, and Writing-review and editing; SW: Data curation, Formal analysis, Investigation, Software, Validation, Visualization, and Writing-review and editing; HO: Conceptualization, Investigation, Methodology, Resources, Validation, and Writing-review and editing.

## References

Anderson, T. J., & MacAskill, M. R. (2013). Eye movements in patients with neurodegenerative disorders. Nature Reviews Neurology, 9(2), 74–85.

Armstrong, T., & Olatunji, B. O. (2012). Eye tracking of attention in the affective disorders: A meta-analytic review and synthesis. Clinical psychology review, 32(8), 704–723.

Benfatto, M. N., Seimyr, G.Ö., Ygge, J., Pansell, T., Rydberg, A., & Jacobson, C. (2016). Screening for Dyslexia Using Eye Tracking During Reading (Version 1). figshare. 10.6084/m9.figshare.c.3521379.v1

Carlsson, G. (2014). Topological pattern recognition for point cloud data. Acta Numerica, 23, 289–368.

Chambers, E. W., De Silva, V., Erickson, J., & Ghrist, R. (2010). Vietoris–rips complexes of planar point sets. Discrete & Computational Geometry, 44(1), 75–90.

Deng, S., Prasse, P., Reich, D. R., Dziemian, S., Stegenwallner-Schütz, M., Krakowczyk, D., … & Jäger, L. A. (2022, September). Detection of ADHD based on eye movements during natural viewing. In Joint European Conference on Machine Learning and Knowledge Discovery in Databases (pp. 403–418). Cham: Springer Nature Switzerland.

Fernández, G., Mandolesi, P., Rotstein, N. P., Colombo, O., Agamennoni, O., & Politi, L. E. (2013). Eye movement alterations during reading in patients with early Alzheimer disease. Investigative ophthalmology & visual science, 54(13), 8345–8352.

Fletcher, W. A., & Sharpe, J. A. (1986). Saccadic eye movement dysfunction in Alzheimer’s disease. Annals of Neurology: Official Journal of the American Neurological Association and the Child Neurology Society, 20(4), 464–471.

Goto, Y., Hatakeyama, K., Kitama, T., Sato, Y., Kanemura, H., Aoyagi, K., … & Aihara, M. (2010). Saccade eye movements as a quantitative measure of frontostriatal network in children with ADHD. Brain and Development, 32(5), 347–355.

Jang, H., Soroski, T., Rizzo, M., Barral, O., Harisinghani, A., Newton-Mason, S., … & Field, T. S. (2021). Classification of Alzheimer’s disease leveraging multi-task machine learning analysis of speech and eye-movement data. Frontiers in Human Neuroscience, 15, 716670.

Kuskowski, M. A. (2013). Eye movements in progressive cerebral neurological disease. In Neuropsychology of Eye Movement (pp. 147–176). Psychology Press.

Lee, D. Y., Shin, Y., Park, R. W., Cho, S. M., Han, S., Yoon, C., … & Kim, S. J. (2023). Use of eye tracking to improve the identification of attention-deficit/hyperactivity disorder in children. Scientific Reports, 13(1), 14469.

Leigh, R. J., & Zee, D. S. (2015). The neurology of eye movements. Oxford University Press, USA.

Lueck, K. L., Mendez, M. F., & Perryman, K. M. (2000). Eye movement abnormalities during reading in patients with Alzheimer disease. Cognitive and Behavioral Neurology, 13(2), 77–82.

Meng, F., Li, F., Wu, S., Yang, T., Xiao, Z., Zhang, Y., … & Luo, X. (2023). Machine learning-based early diagnosis of autism according to eye movements of real and artificial faces scanning. Frontiers in Neuroscience, 17, 1170951.

Munoz, D. P., Armstrong, I. T., Hampton, K. A., & Moore, K. D. (2003). Altered control of visual fixation and saccadic eye movements in attention-deficit hyperactivity disorder. Journal of neurophysiology, 90(1), 503–514.

Munoz, D. P., Hampton, K. A., Moore, K. D., & Goldring, J. E. (1999). Control of purposive saccadic eye movements and visual fixation in children with attention-deficit hyperactivity disorder. In Current Oculomotor Research: Physiological and Psychological Aspects (pp. 415–423). Boston, MA: Springer US.

Nilsson Benfatto, M., Öqvist Seimyr, G., Ygge, J., Pansell, T., Rydberg, A., & Jacobson, C. (2016). Screening for dyslexia using eye tracking during reading. PloS one, 11(12), e0165508.

Parkinson, J., & Maxner, C. (2005). Eye movement abnormalities in Alzheimer disease: case presentation and literature review. American Orthoptic Journal, 55(1), 90–96.

Przybyszewski, A. W., Kon, M., Szlufik, S., Dutkiewicz, J., Habela, P., & Koziorowski, D. M. (2014). Data mining and machine learning on the basis from reflexive eye movements can predict symptom development in individual Parkinson’s patients. In Nature-Inspired Computation and Machine Learning: 13th Mexican International Conference on Artificial Intelligence, MICAI 2014, Tuxtla Gutiérrez, Mexico, November 16-22, 2014. Proceedings, Part II 13 (pp. 499–509). Springer International Publishing.

Raatikainen, P., Hautala, J., Loberg, O., Kärkkäinen, T., Leppänen, P., & Nieminen, P. (2021). Detection of developmental dyslexia with machine learning using eye movement data. Array, 12, 100087.

Robins, V. (1999). Towards computing homology from finite approximations. In Topology proceedings (Vol. 24, No. 1, pp. 503–532).

Rojas-Líbano, D., Wainstein, G., & Ossandon, T. (2019). Pupil Size, Eye-tracking and Neuropsychological Dataset from ADHD-diagnosed and control participants performing a cognitive task. (Version 3). figshare. 10.6084/m9.figshare.7218725.v3

Rojas-Líbano, D., Wainstein, G., Carrasco, X., Aboitiz, F., Crossley, N., & Ossandón, T. (2019). A pupil size, eye-tracking and neuropsychological dataset from ADHD children during a cognitive task. Scientific data, 6(1), 25.

Rommelse, N. N., Van der Stigchel, S., & Sergeant, J. A. (2008). A review on eye movement studies in childhood and adolescent psychiatry. Brain and cognition, 68(3), 391–414.

Tauzin, G., Lupo, U., Tunstall, L., Pérez, J. B., Caorsi, M., Medina-Mardones, A. M., … & Hess, K. (2021). giotto-tda:: A topological data analysis toolkit for machine learning and data exploration. Journal of Machine Learning Research, 22(39), 1–6.

Tseng, P. H., Cameron, I. G., Pari, G., Reynolds, J. N., Munoz, D. P., & Itti, L. (2013). High-throughput classification of clinical populations from natural viewing eye movements. Journal of neurology, 260, 275–284.

Vabalas, A., Gowen, E., Poliakoff, E., & Casson, A. J. (2020). Applying machine learning to kinematic and eye movement features of a movement imitation task to predict autism diagnosis. Scientific reports, 10(1), 8346.

Vietoris, L. (1927). Über den höheren Zusammenhang kompakter Räume und eine Klasse von zusammenhangstreuen Abbildungen. Mathematische Annalen, 97(1), 454–472.

Yoo, J. H., Kang, C., Lim, J. S., Wang, B., Choi, C. H., Hwang, H., … & Kim, J. W. (2024). Development of an innovative approach using portable eye tracking to assist ADHD screening: a machine learning study. Frontiers in Psychiatry, 15, 1337595.

Zemblys, R., Niehorster, D. C., Komogortsev, O., & Holmqvist, K. (2018). Using machine learning to detect events in eye-tracking data. Behavior research methods, 50, 160–181.

